# Hyperbolic matrix factorization reaffirms the negative curvature of the native biological space

**DOI:** 10.1101/2020.12.21.423806

**Authors:** Aleksandar Poleksic

## Abstract

Past research in systems biology has taken for granted the Euclidean geometry of biological space. This has not only drawn parallels to other fields but has also been convenient due to the ample statistical and numerical optimization tools available to address the core task and downstream machine learning problems. However, emerging theoretical studies now demonstrate that biological databases exhibit hierarchical topology, characterized by heterogeneous degree distribution and a high degree of clustering, thus contradicting the flat geometry assumption. Namely, since the number of nodes in hierarchical structures grows exponentially with node depth, the biological networks naturally reside in a hyperbolic space where the circle circumference and disk area are the exponential functions of the radius. To test these claims and assess potential benefits of the applications grounded in the above hypothesis, we have developed a mathematical framework and an accompanying computational procedure for matrix factorization and implied biological relationship inference in hyperbolic space. Not only does our study demonstrate a significant increase in the accuracy of hyperbolic embedding compared to Euclidean embedding, but it also shows that the latent dimension of an optimal hyperbolic embedding is by more than an order of magnitude smaller than the latent dimension of an optimal Euclidean embedding. We see this as additional evidence that hyperbolic geometry, rather than Euclidean, underlines the biological system.

## Introduction

Although biological databases have seemingly high dimensionality, their intrinsic dimension is relatively low. For instance, many genes in a genomic database are co-expressed and a significant portion of variations in these databases can be explained by a few variables, including the cell type, the number of detected transcripts, or a gene program [1]. In a different example, the low intrinsic dimension of databases of adverse drug reactions is due to associations of side-effects to particular chemical substructures and their combinations. In other words, it is well established that drugs sharing chemical substructures are likely to give rise to the same side-effects [2,3]. Thus, representing these and other biomedical entities as points in a low dimensional space, in a way that preserves distances between those entities, is essential for many systems biology tasks, including classification, clustering, link prediction, and data visualization.

Matrix factorization is the method of choice for dimensionality reduction and relationship inference [4–21]. It is the key technique used to predict drug-disease associations, drug-target interactions, drug-side effect associations, and disease-associated genes, to name a few. Given a matrix *R_m×n_* of relationships between the elements from two biological domains *A* and *B,* the goal is to find the best approximation *R* ≈ *UV^T^* as the product of two lower dimensional matrices *U_m×d_* and *V_n×d_.* The rows of *U* and *V* can be seen as points in ℝ^*d*^ and are called the *feature* or *latent* representations of elements from *A* and *B,* respectively, while the entries of *UV^T^* can be thought of as the predicted association scores between those elements. In a different interpretation, if we view the elements of *A* and *B* as the network nodes and elements of *R* as the network edges, then the rows of *U* and *V* represent the solutions of the heterogeneous network embedding problem, namely, the problem of embedding a simple two-domain network.

The research on matrix factorization and network embedding has historically focused on computational and statistical aspects of the particular problem of interest, taking the Euclidean geometry of the latent space for granted. However, studies have recently emerged suggesting that the native space of complex networks is not the Euclidean space ℝ^*d*^, but the hyperbolic space of negative Gaussian curvature [22–28]. A compelling argument in support of this hypothesis is that complex networks resemble hierarchical, tree-like organization [29] and that discrete hyperbolic geometry is the geometry of trees. Several filed specific applications have also been developed, demonstrating the benefits of data representation in a negatively curved space [1,30–37].

Just like the Internet and social networks [22,38], biological networks are organized into many small topological modules, grouped together in a hierarchical manner to form a hierarchical network [25,39]. Embedding such a network in a flat Euclidean space leads to distortion in spatial arrangement of the feature vectors representing individual biological entities. In contrast, the hyperbolic space can accommodate exponential growth of the number of relevant network features since the circumference of a hyperbolic circle and the disk area are exponential functions of the radius. Nevertheless, in spite of this simple, convincing argument, the above assumption on the nature of the biological space has never been tested in practice nor it has been built upon to create practical and useful biological applications.

In this study we lay out the theoretical foundation for biological native space representation and develop the corresponding computational procedure for hyperbolic matrix factorization. More specifically, we take advantage of recent advances in the development of probabilistic models and numerical optimization, including the work on pseudo-hyperbolic Gaussian distribution [40] and a new technique for gradient descent in hyperbolic space [41,42] to learn the latent node representation and, in turn, compute probabilities of associations between biological entities. Our benchmarking data clearly and consistently demonstrates a significant advantage in accuracy and a drastic reduction in latent space dimensionality of hyperbolic matrix factorization.

## Results

We carried out a head-to-head comparison of the Euclidean and hyperbolic matrix factorization using the probabilistic logistic matrix factorization framework. Originally proposed for item recommendations in the media industry [43,44], logistic matrix factorization has become the state-of-the-art method for biological relationship inference [45–53]. Given an incomplete binary matrix *R* = (*r_i, j_*) of observed relationships between elements of two domains *A* and *B*, the probability of *a^i^* ∈ *A* and *b^j^* ∈ *B* being related (ry = 1) is modeled as

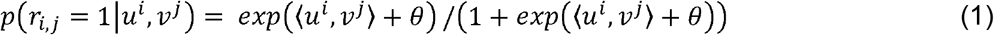

where *u^i^* ∈ ℝ^*d*^ and *v^i^* ∈ ℝ^*d*^ are the latent vector representations of *a^i^* and *b^j^*, respectively, 〈·,·〉 represents the Euclidean inner product, and *θ* is a trainable parameter [44]. The latent vectors are learned through the maximum a posteriori estimation using the alternating gradient descent technique.

We developed a hyperbolic variant of the above method by considering *u^i^* and *v^i^* as the points in *d*-dimensional hyperbolic space 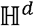 and by replacing the Euclidean inner product 〈*u^i^*, *v^j^* 〉 in (1) by the Lorentzian product 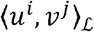 (details in Methods section).

We trained the parameters of each method (Euclidean and hyperbolic) separately on each benchmarking data set, using the gradient descent in hyperbolic space and an extensive grid search. Table 1 shows the comparison in accuracy of Euclidean and hyperbolic factorization, as measured by the AUC and the AUPR scores on seven biological databases: drug-target interaction databases Nr, Gpcr, Ion, Enz, [54], and DrugBank [55], drug-disease association database PharmacoDB [56], and the SIDER database of adverse drug reactions [57]. Fig. 1 shows the latent dimension of an optimal Euclidean factorization as well as the smallest hyperbolic dimension that yields better or equal AUC and AUPR scores.

**Table 1.** Comparison of “bare-bones” (no side information) hyperbolic and Euclidean matrix factorization using 10 rounds of 5-fold cross validation.

|  | AUPR |  | AUC |  |
| --- | --- | --- | --- | --- |
|  | Hyperbolic | Euclidean | Hyperbolic | Euclidean |
| **Nr** | **0.48(0.05)** | 0.35(0.00) | **0.84(0.01)** | 0.76(0.01) |
| **Gpcr** | **0.63(0.00)** | 0.59(0.01) | **0.91(0.00)** | 0.87(0.01) |
| **Ion** | **0.87(0.00)** | 0.84(0.00) | **0.97(0.00)** | 0.97(0.00) |
| **Enzyme** | **0.82(0.00)** | 0.74(0.00) | **0.97(0.00)** | 0.95(0.00) |
| **DrugBank** | **0.52(0.00)** | 0.44(0.00) | **0.95(0.00)** | 0.92(0.00) |
| **PharmDB** | **0.35(0.00)** | 0.27(0.01) | **0.88(0.00)** | 0.78(0.00) |
| **Sider** | **0.50(0.05)** | 0.49(0.00) | **0.95(0.00)** | 0.94(0.00) |

**Fig. 1.**
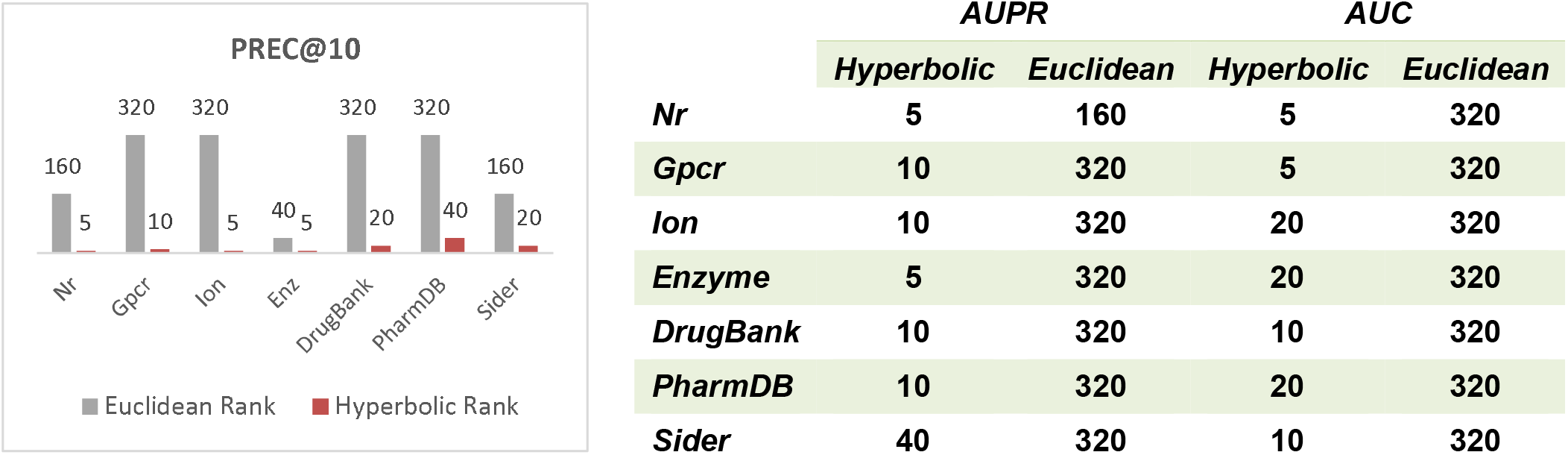
Optimal Euclidean rank (latent space dimension) and the hyperbolic rank yielding the same or better classification scores with respect to PREC@10 metric (left) and AUC and AUPR (right).

As seen in table 1 and Fig. 1., the hyperbolic matrix factorization routinely outperforms the Euclidean factorization in every benchmark and across fundamentally different classification measures. Moreover, the hyperbolic matrix factorization achieves superior accuracy at latent dimensions that are by an order of magnitude smaller compared to dimensions needed for optimal Euclidean embedding. Specifically, optimal Euclidean factorization is most often achieved at ranks exceeding 300. In contrast, hyperbolic factorization needs only 5 or 10 latent features to achieve the same or better classification scores.

We obtained similar results on MovieLens [58] and Epinions [59] databases (data not shown). While our focus is on biomedical applications, the benchmarking data on movie rating and social networking datasets further confirms the hyperbolic nature of the geometry of complex networks and superiority of hyperbolic factorization.

## Methods

Hyperbolic geometry can be modeled on the *n*-dimensional hyperboloid in the Lorentzian space ℝ^*n*,1^, where ℝ^*n*,1^ is a copy of ℝ^*n*+1^ equipped with a bilinear form 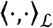 defined as

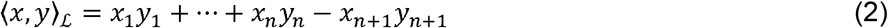

Hyperbolic space is represented by one sheet of the two-sheeted hyperboloid

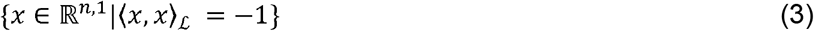

(which can be thought of as a sphere of radius 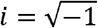) namely,

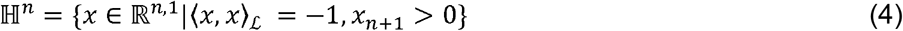

It can be shown that the bilinear form 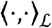 restricted on the tangent space at a point *p* ∈ 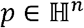, defined by

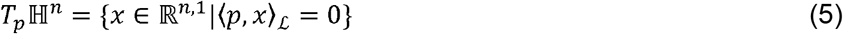

is positive definite, thereby representing a genuine Riemannian metric on 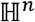. The distance between two points *x,* y ∈ 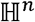 is given by

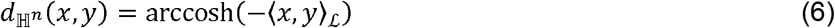

An interesting and in the biological context insightful property of the hyperbolic space is that the shortest path between two random points in 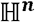 that are far away from the vertex ***μ_0_*** as almost the same length as the path through the vertex (Fig. 2). This resembles the property of the distance function on trees, where the shortest path between two randomly selected nodes deep in the tree is almost of the same length as the path through the root. A good biological analogue is a phylogenetic tree, where the shortest path between two arbitrary selected organisms is almost the path through their most ancient ancestor (root).

**Fig. 2.**
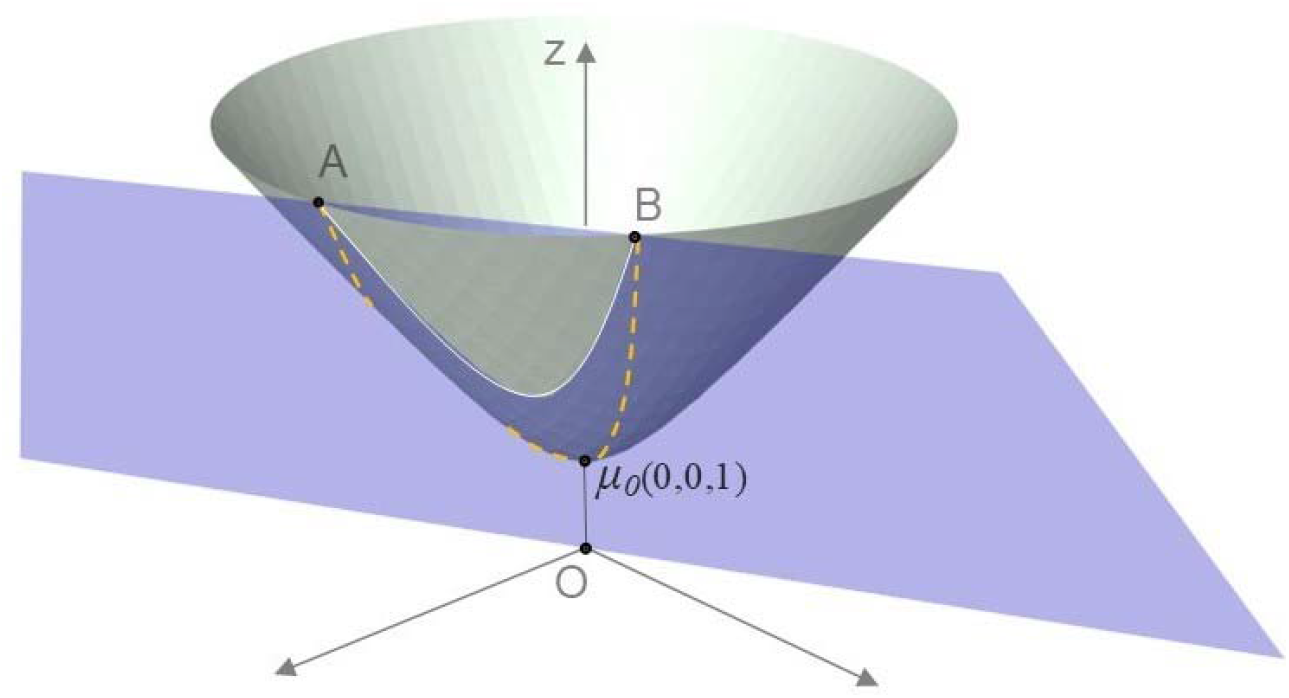
Hyperboloid model of ℍ^2^ The shortest path between points A and B is the line of intersection of the hyperboloid with the plane (blue) through A, B and O. As A and B are moving away from the vertex *μ*_0_, the length of this geodesic line (white) is almost the same as the length of the path through *μ*_0_ (orange).

### The theoretical foundation

While the general theory, presented below, can be tailored to different matrix factorization techniques (or, at least to those in which the objective function is a function of the hyperbolic inner product), we illustrate it in the framework of logistic matrix factorization. Our choice is due to the explicit statistical formulation of logistic matrix factorization, simplicity of presentation, and its high accuracy in biological applications [47–53].

Let *A* and *B* be two biological domains consisting of entities (nodes) 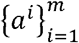 and 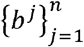 respectively. Denote by *R* =(*r_i,j_*)_*m*×*n*_ the matrix of relationships (edges) between the elements of *A* and *B*. Specifically, *r_ij_* = 1 if *a^i^* is related to *b^j^* and *r_i, j_* = 0 otherwise (no relationship or unknown). Let 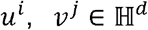 be the latent vector representations of *a^i^* and *b^j^*, respectively, where *d* ≪ max (*m,n*). Denote by *e_j,j_* the event that *a^i^* and are related. Following the Euclidean case, we model the probability of *e_i,j_* as the logistic function in the Lorentz space ℝ^*d*,1^

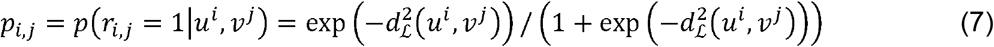

where 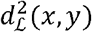 denotes the squared *Lorentzian distance* [60] between the points *x,y* ∈ ℍ^*d*^, namely

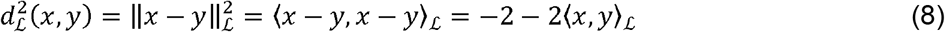

Denote by *W* = (*w_i,j_*)_*m*×*n*_ our confidence in the entries *r_i,j_* of the interaction matrix *R*. In many practical applications, *w_i,j_* = 1 if and *w_i,j_* = *c* if *r_i,j_* = 1, where *c* > 1 is a constant. In general, the idea is to assign higher weights to trustworthy pairs i.e., the pairs of biological concepts (*a^i^*, *b^j^*) for which we have higher confidence of association. Given the weights *w_i,j_*, the likelihood of *r_i,j_* given *u^i^* and *v^i^* is

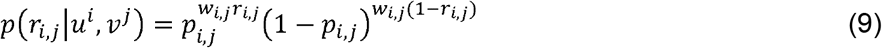

Assuming the independence of events *e_i,j_*, it follows that

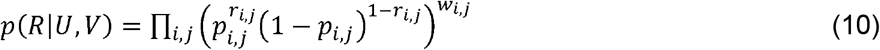

where *U* and *V* represent the matrices of latent preferences of elements from *A* and *B,* respectively. In other words, the *i^th^* row of *U* is the vector *u^i^* and *i^th^* row of *V* is *v^i^*.

### Computing the prior distribution

Our goal is to derive the probability *p*(*U,V*|*R*) from (10) through the Bayesian inference, which requires placing a prior on latent vectors (rows of *U* and *V*).

Borrowing from the recent work on wrapped normal distribution in hyperbolic space [40], we define the prior distributions

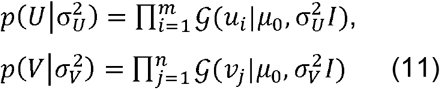

where 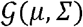 is the pseudo-hyperbolic Gaussian distribution and *μ*_0_ = (0,…, 0,1) is the vertex of the hyperboloid i.e. the origin of the hyperbolic space.

The pseudo-hyperbolic Gaussian distribution extends the notion of Gaussian distribution on the hyperbolic space (Fig. 3). In short, for 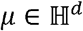 and positive definite Σ, sampling from 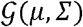 can be thought of as a three step process: (a) sample a vector 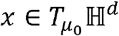from 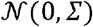 (b) transport *x* along the geodesic joining the points 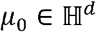 and 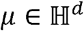 to 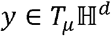, (c) project *y* to 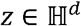.

**Fig. 3.**
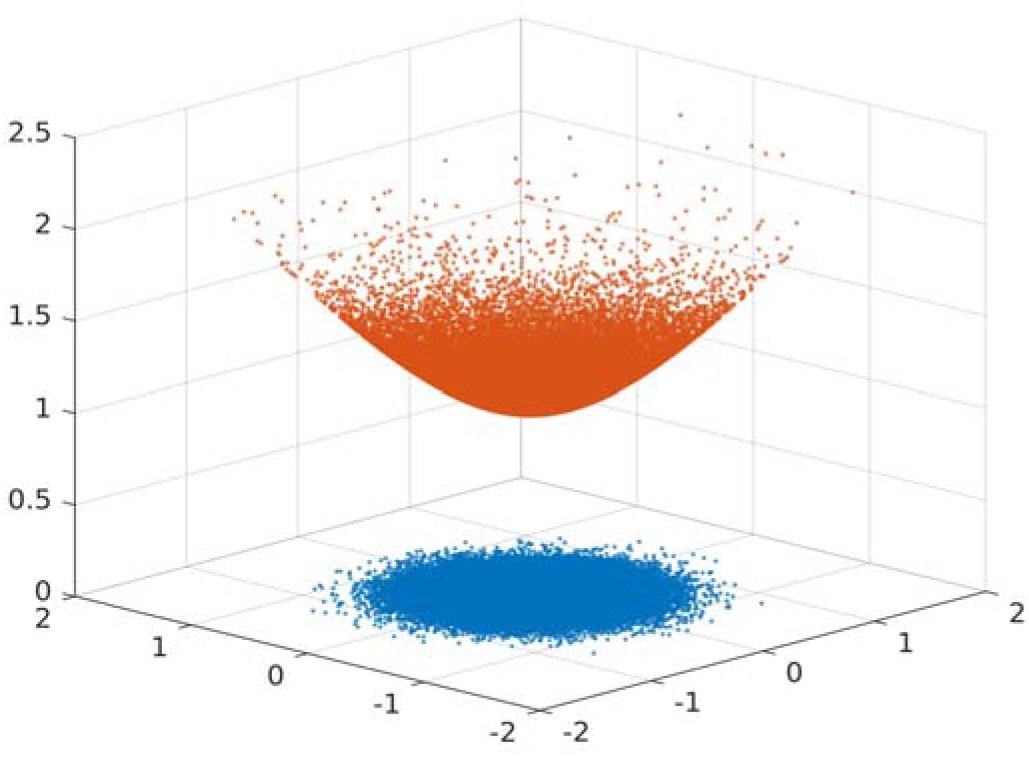
100,000 samples from 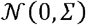 (blue) and the corresponding samples from 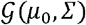 (red), Σ = 0.1 · *I*_2×2_

The step (b) is carried out using the parallel transport 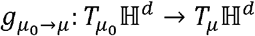 (Fig. 4):

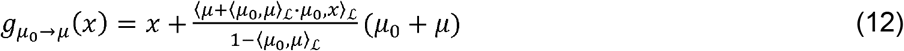

while the step (c) uses the exponential map 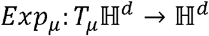 (Fig. 4), defined by

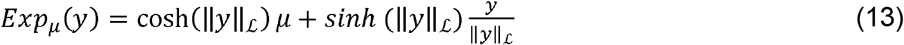

where 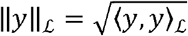.

**Fig. 4.**
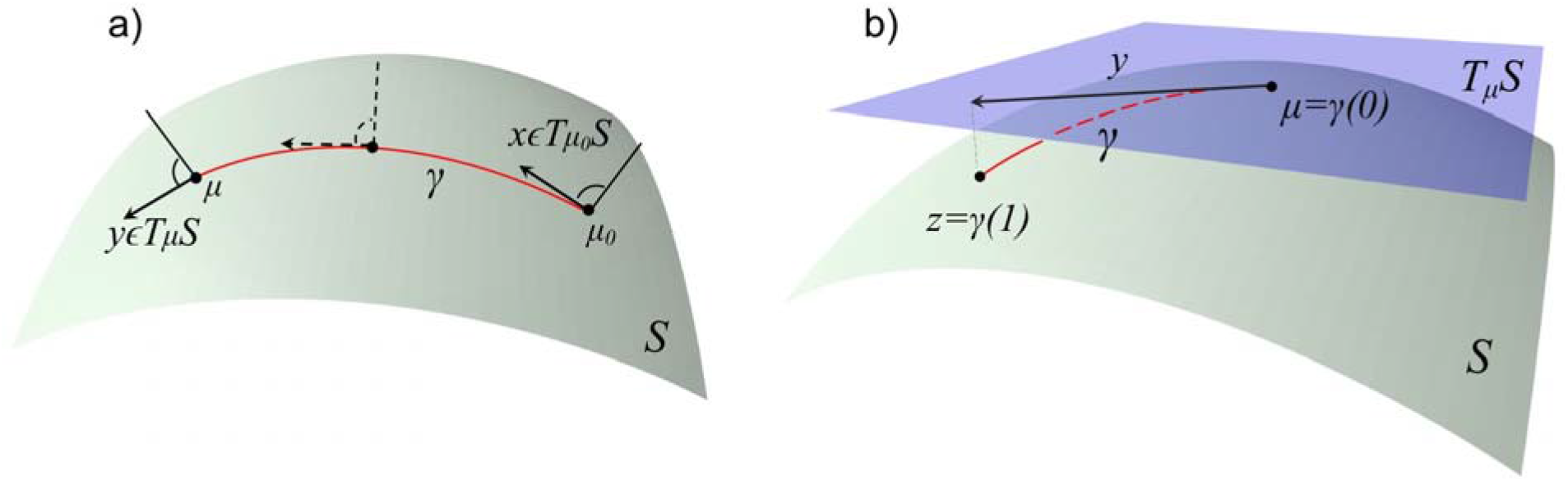
(a) Parallel transport of *x* ∈ *T*_*μ*0_*S* to *y* ∈ *T_μ_S* along the geodesic *γ*, where *μ*_0_ = *γ*(0) and *μ* = *γ*(1). (b) The exponential map.

It is not difficult to show that the length of the geodesic joining *μ* to *Exp_μ_(*y*)* on 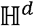 is equal to 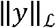, i.e., 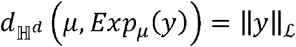. The relationship between the probability densities 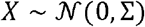 and 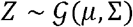 is

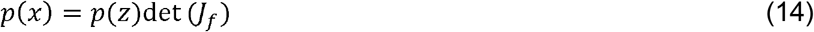

where *f* = *Exp_μ_* ° *g*_*μ*_0_*→μ*_ and det(*J_f_*) denotes the determinant of the Jacobian 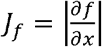 [40]. It can also be shown that

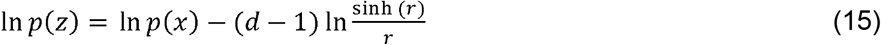

where 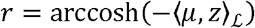 [40].

### The loss function

With the prior placed on *U* and *V,* we return to calculating the posterior probability *p*(*U, V*|*R*) through the Bayesian inference:

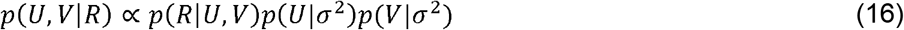

Taking the negative logarithm of the posterior distribution (16), we arrive at the closed form expression for the loss function

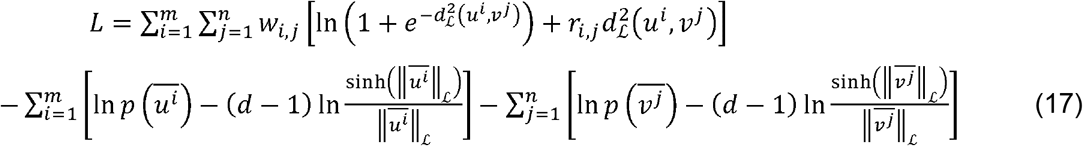

In the expression above, *p* is the probability density function of the normal distribution 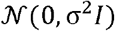 in the tangent space 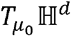 at the vertex *μ*_0_ = (0,…,0,1). For 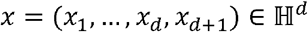

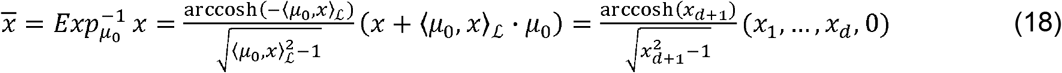

It follows that

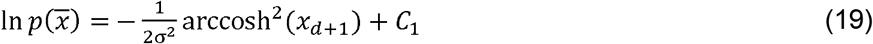

where *C*_1_ is a constant. Since

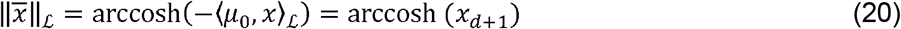

we have

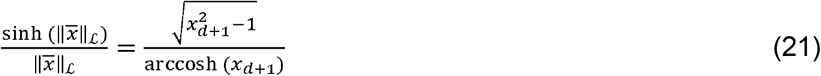

Hence, our loss function will have the form:

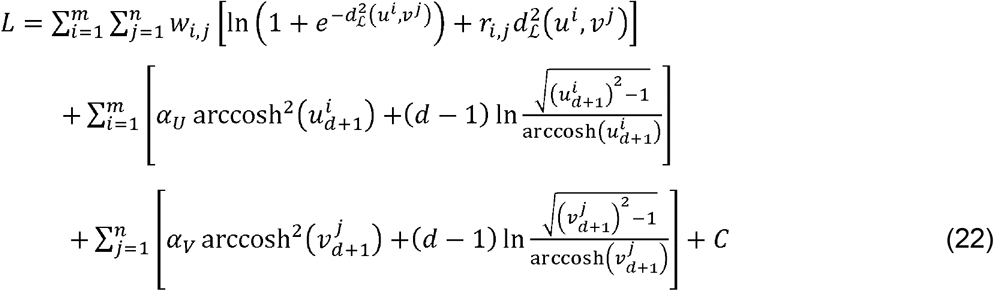

where 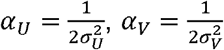 are trainable parameters and *C* is a constant.

### Alternating gradient descent in hyperbolic space

Finding a point of (local) minimum of a real function defined in a d-dimensional Euclidean space ℝ^*d*^ is routinely accomplished using the gradient descent. We adopt a similar technique for finding the point 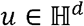 of a local minimum of any real valued function 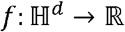 [41]. For this strategy to work, the function *f* has to be defined is in the ambient space ℝ^*d*,1^ of 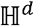, as well as on 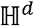. Specifically, given the initial value *u* = *u*^(0)^ and a step size *η*, the gradient descent in hyperbolic space can be carried out by repeating the following steps:

1. Compute the gradient 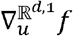
2. Project 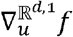 orthogonally to vector 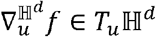
3. Set 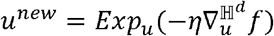

The gradient 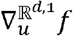 of *f* in the ambient space ℝ^*d*,1^ is a vector of partial derivatives

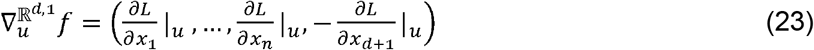

The representation above (note the negative sign of the last vector’s component) follows directly from the definition of the gradient:

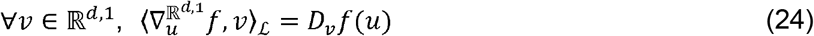

The orthogonal projection from the ambient onto the tangent space in (step 2 above) is given by

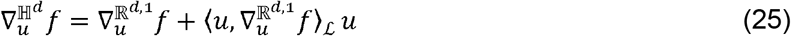

We will use the “alternating gradient descent” method to minimize the error function *L_A,B_* given in (22). The partial derivatives of *L_A,B_* are relatively easy to compute:

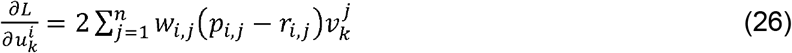

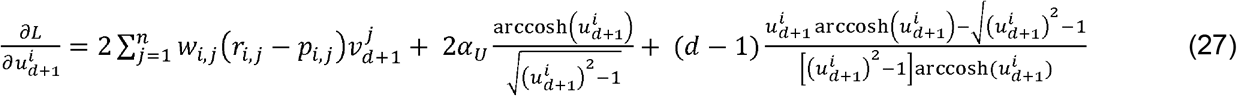

**ALGORITHM.**
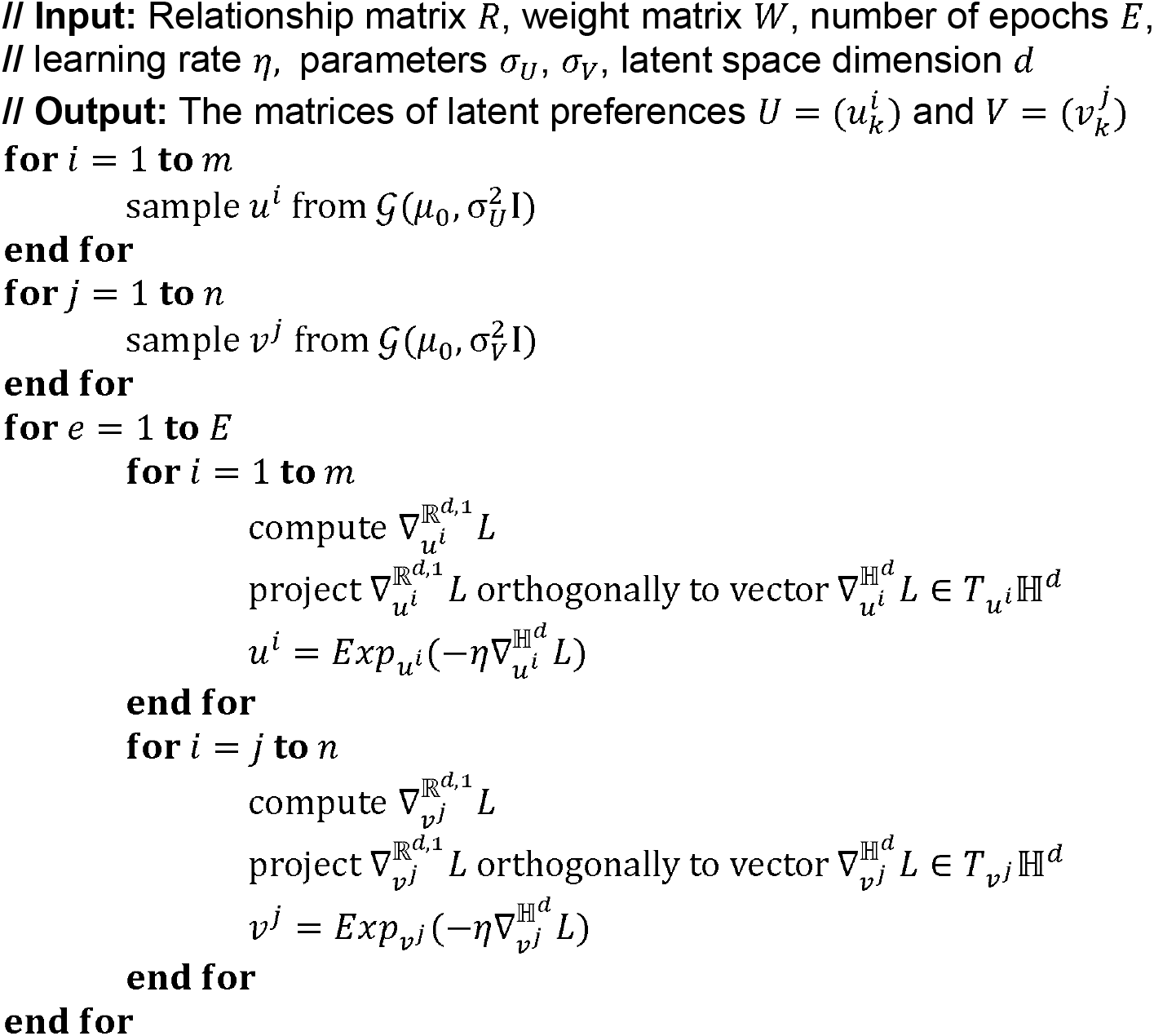
Hyperbolic Alternating Gradient Descent

### Generalization to spaces of arbitrary curvature and field specific applications

There are several directions for future research that can enhance the utility of hyperbolic matrix factorization. One approach is to generalize the theory to spaces of arbitrary negative curvature. The hyperbolic space of curvature −1/*β*, where *β* > 0, is defined as

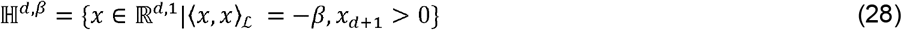

Curvature specific matrix factorization can be carried out by generalizing (7) to

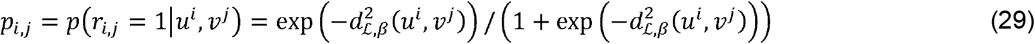

where 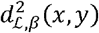 denotes the generalized Lorentzian distance [60] between the points 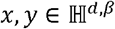:

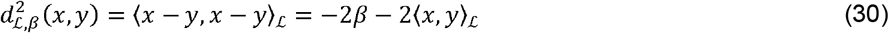

However, extending the factorization procedure to 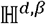 requires the notion of pseudo-hyperbolic Gaussian distribution from 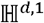 to 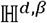.

In certain applications, such as recommender systems, the matrix factorization is supplemented with other techniques to improve the accuracy of relationship inference. In one approach, the *neighborhood regularization* is used to ensure that similar entities from the domain *A* are in relationship with similar entities from the domain *B.* The existing techniques for incorporating side information [45,46,47,61] can be carried out in 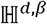 by adding

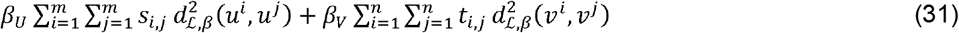

to the loss function *L,* where *β*_U_ *β_v_* are trainable parameters.

Similarly, the arrival of a new entity in one of the domains, coined as the *cold-start* problem, can be addressed with the hyperbolic variant of the weighted-profile method [45,46,62]. In Euclidean case, the latent representation *u^i^* ∈ ∝^*d*^ of the element *a^i^* ∈ *A* that is not in an observed relationship with any *b^j^* ∈ *B* is often computed as the weighted combination of the rows *u*^j^ ∈ *U* most similar to *u^i^*:

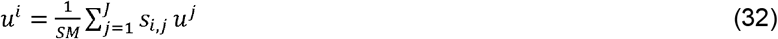

where 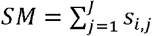. We note that the point *u^i^* in (28) represents the center of mass of the points 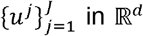. Specifically, *u^i^* is the solution to the minimization problem

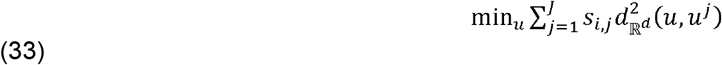

where 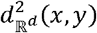 is the squared Euclidean distance between the points *x* and *y.*

The corresponding hyperbolic weighted-profile method can utilize a closed form solution for the center of mass in the hyperbolic space, i.e. the solution of the minimization problem

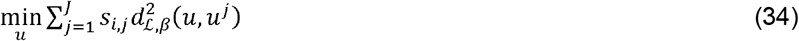

given recently by Law *et al.* [63], namely

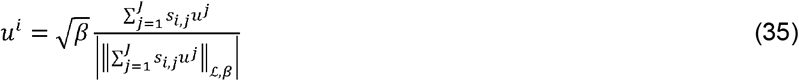

where 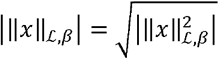.

## Conclusion

Matrix factorization is one of the main techniques used in systems biology for predicting the relationships between the elements from a pair of biological domains. The key idea is to complete an input matrix of observed relationships using its close low-rank approximation i.e., the product of two lower dimensional matrices. The technique results in the representation of biological objects as points in a low dimensional Euclidean space. This latent space representation is essential in nearly all downstream machine learning tasks, such as classification, clustering, visualization, and the overall understanding of the structure and dynamics of a biological system.

Past research in systems biology has taken the Euclidean geometry of the biological space for granted. This has been convenient due to the availability of advanced analytic, numerical, statistical and machine learning procedures in the Euclidean space. However, recent theoretical studies suggest that the hyperbolic geometry, rather than Euclidean, underpins all complex networks in general and biological networks in particular. Therefore, a radical shift in data representation is necessary to obtain an undistorted view of the biological space and, in turn, ensure further progress in systems biology and related fields.

We have developed and benchmarked a technique for a probabilistic hyperbolic matrix factorization and resulting hyperbolic space embedding. We demonstrate that the Lorentzian model of hyperbolic space allows for a closed form expression of the key transformations and techniques required for latent space dimensionality reduction. Our method builds upon some recent advances in the development of probabilistic models and numerical optimization in hyperbolic space to learn an optimal hyperbolic embedding and to compute the probabilities of relationships between biological entities. The benchmarking tests performed on different types of biomedical data clearly demonstrate a significant increase in accuracy and a drastic reduction in latent space dimensionality of hyperbolic embedding compared to Euclidean embedding. These findings confirm the negative curvature of the native biological space.

The techniques presented in this study can be generalized to hyperbolic spaces of any given curvature. Finding the proper curvature of the space that underlines a particular biological database will result in a more accurate latent representation and, in turn, more accurate biological relationship prediction.

